# Interoceptive autonomic regulation in typical development and autism spectrum disorder: A computational model integrating multiple physiological systems

**DOI:** 10.64898/2026.02.19.706737

**Authors:** Ruichen Li, Hao Liu, Yukie Nagai

## Abstract

**Background:** Interoceptive cardiovascular signals, including heart rate (HR) and blood pressure (BP), arise from coordinated sympathetic (SNS) and parasympathetic (PSNS) regulation and contribute to affective and cognitive processes. Although atypical autonomic nervous system (ANS) modulation has been reported in autism spectrum disorder (ASD), the dynamical structure underlying branch-specific coordination remains insufficiently characterized.

**Objective:** To estimate latent ANS regulatory structure in typically developing (TD) and ASD individuals using a computational modeling framework.

**Methods:** A closed-loop computational model integrating cardiovascular, respiratory, and autonomic dynamics was developed. ANS regulation was formalized using three autonomic control modes (coupled reciprocal, coupled nonreciprocal, and uncoupled) and parameterized by the relative activity of SNS and PSNS branches. HR and BP responses to the head-up tilt (HUT) test were simulated, and regulatory surfaces were compared with empirical HR and BP data from TD and ASD groups. Additional simulations under normal respiration, deep respiration, and absence of respiration evaluated mean arterial pressure (MAP) regulation across varying SNS–PSNS activity combinations.

**Results:** TD individuals exhibited differentiated SNS–PSNS coordination patterns across control modes, whereas ASD individuals showed convergence of relative SNS–PSNS activity. In TD, HR and BP distributions under coupled reciprocal mode were most consistent with expected physiological responses, characterized by high SNS and low PSNS activity during postural challenge. In ASD, empirical data extended toward regions of relatively higher PSNS weighting. Incorporation of deep respiration enhanced MAP reduction during BP recovery, particularly under over-elevated SNS activity.

**Conclusion:** This study provides a mechanistic, state-space characterization of autonomic coordination in TD and ASD populations, enabling inference of latent autonomic regulation from measurable interoceptive phenotypes and identifying respiration as a model-based regulatory lever that augments cardiovascular stabilization.

## 1. Introduction

The autonomic nervous system (ANS), consisting primarily of the sympathetic and parasympathetic nervous systems (SNS and PSNS), serves as the principal regulatory network governing rhythmic organ functions and modulating interoceptive states to support homeostasis and adaptive responses [1]-[7]. Cardiovascular and respiratory afferents converge in key central regions such as the brainstem, insula, and anterior cingulate cortex, shaping autonomic output and bodily-state perception [8]. Interoceptive rhythms such as arterial pressure and cardiac cycling activate baroreceptors within the aortic arch and carotid sinus, triggering sympathetic–parasympathetic adjustments to maintain circulatory stability [9],[10]. In addition, respiratory activity modulates venous return and stroke volume, generating respiratory sinus arrhythmia that reflects cardiopulmonary coupling [11],[12]. Variations in heart rate (HR) and blood pressure (BP) are therefore commonly used as non-invasive indices for assessing sympathetic–parasympathetic balance [13]-[16].

Growing evidence indicates that atypical interoceptive processing is a hallmark of autism spectrum disorder (ASD), with imprecise interoceptive prediction contributing to sensory atypicality and disrupted emotional-cognitive processing [21]-[25]. Because interoceptive signals are conveyed through autonomic pathways, altered interoceptive processing may originate from atypical autonomic regulation. Heightened ANS dysfunction in ASD individuals have been reported to affect cardiopulmonary and following social functioning [26]-[27]. During anxiety, individuals with ASD also exhibit elevated HR alongside reduced electrodermal activity and skin temperature, suggesting atypical SNS–PSNS coordination that may serve as a physiological marker of dysregulation [28]. However, contrasting findings have also emerged that presented typical baseline parasympathetic activity and normal autonomic responses to social stimuli in many autistic children [29], while insufficient sympathetic activation and low physiological arousal were also found in ASD [30], indicating a contrasting profile to excessive sympathetic dominance reported elsewhere. Consequently, the inconsistency in empirical findings on autonomic dynamics constrains the ability to identify differences of interoceptive modulation between typically developing (TD) and ASD individuals, which need for a quantitative and mechanistic framework capable of identifying how differences in autonomic function shape interoceptive cardiovascular responses.

Predictive coding and active inference models conceptualize interoception as generative inference over visceral states, governed by predictions, prediction errors, and precision weighting. Computational formulations reviewed by Petzschner et al. [53] and implemented by Allen et al. [54] enable parameterized characterization of interoceptive regulation, offering a principled framework for comparing interoceptive processing across individuals and populations. However, the existing models focus on a single physiological system or a limited set of interoceptive channels and remain limited in their capacity to capture coordinated regulation across multiple physiological systems. Moreover, their sensitivity to TD-ASD differences has not yet been systematically evaluated. These limitations constrain the explanatory power of current interoceptive models in accounting for differences in interoceptive regulation between TD and ASD populations and highlight the need for computational approaches that integrate multiple physiological systems while providing greater resolution for characterizing group-specific regulatory mechanisms. The objectives of this study are to establish a computational model integrating multiple physiological systems including cardiovascular, respiratory, and autonomic nervous systems. By analyzing the time-varying HR and BP variations, this computational model enables to clarify how ANS functions give rise to differences in HR and BP across TD and ASD populations, and to evaluate whether self-regulation strategies that modulate autonomic output promote recovery of impaired interoceptive cardiovascular rhythms. Achieving these objectives advances mechanistic insight into autonomic contributions to interoceptive regulation and provides a computational foundation for personalized physiological intervention strategies.

## 2. Related works

### 2.1 Autonomic regulation strategy

Traditionally, autonomic regulation of interoceptive signals has been conceptualized as a reciprocal process, in which SNS and PSNS activities change in a mutually antagonistic manner. Berntson et al. [17] extended this view by introducing the concept of autonomic control modes to characterize a broader repertoire of SNS–PSNS coordination patterns. Specifically, autonomic regulation may operate in (1) a coupled reciprocal mode, in which SNS and PSNS activities change in opposing directions; (2) a coupled nonreciprocal mode, in which the two branches are either coactivated or coinhibited; and (3) an uncoupled mode, in which activity in one branch varies independently of the other. Research on hypoxic challenges by Fukuda et al. [18] and Kollai et al. [19] further supports this framework: severe hypoxia elicits reciprocal ANS control—SNS activation accompanied by PSNS withdrawal—to increase HR and BP for oxygen delivery, whereas mild hypoxia induces SNS–PSNS coactivation that increases vascular resistance while stabilizing HR. Existing evidence [20] suggests that flexibility in ANS control modes likely extends across a range of physiological stimuli and psychological states; however, a systematic characterization of how distinct control modes are recruited across contexts remains lacking.

### 2.2 Physiological modeling

Multiscale cardiovascular modeling frameworks have been strategically established and applied based on hemodynamic mechanisms and electrical circuit analogies, ranging from 0D lumped-parameter models for global circulation dynamics, to 1D whole-body arterial network models capturing pulse wave propagation, and 3D regional models resolving local blood flow and vessel biomechanics [31]-[39]. The computational model incorporating respiratory system influences was developed to quantify the effects of different breathing patterns, including normal and deep respiration, on systemic hemodynamics [40]-[41]. Respiratory-induced oscillations were introduced through a dynamic intrathoracic pressure (ITP) function embedded in BP waveforms, representing thoracic pressure fluctuations and revealing characteristic phase coupling between respiration and cardiovascular dynamics. In addition, autonomic control modeling has evolved from early formulations emphasizing isolated baroreflex or cardiovascular mechanics to more integrative frameworks that incorporate bidirectional sympathetic–parasympathetic pathways, gas exchange regulation, and afferent–efferent coupling mechanisms [42]-[44]. The present study defines autonomic function using two dimensions: autonomic control modes [17], and the relative activity levels of the sympathetic and parasympathetic branches. The computational model is initialized using previously reported experimental HR and BP data from TD and ASD groups under external stimulation [45], allowing individualized physiological states to be embedded in the initialization process. Model parameters are subsequently fitted to the observed dynamics to generate analytical solutions that characterize state-dependent patterns of autonomic regulation. The fitted responses are evaluated using three-dimensional interoceptive response surfaces to quantify ANS regulatory behavior in terms of control modes and sympathetic–parasympathetic balance.

## 3. Methods

A closed-loop computational model was developed to simulate the dynamic modulation of HR and BP through autonomic regulation by integrating the cardiovascular system (CVS), respiratory system (RS), and autonomic nervous system (ANS). The overall model architecture is shown by the left panel in Fig 1(a). The CVS submodule generates baseline HR and cyclic BP waveforms, which are subsequently modulated by the ANS submodule under different external stimuli and autonomic control modes. The RS submodule influences the CVS-generated BP waveforms in a unidirectional manner by specifying changes in intrathoracic pressure (ITP) associated with different breathing patterns. Building on existing autonomic regulation models, the present study further formalizes ANS function by explicitly decomposing it into two complementary components: autonomic control modes and the relative activity of the SNS and PSNS branches. The autonomic control modes characterize how SNS and PSNS interact to regulate cardiovascular dynamics, whereas relative activity captures the branch-specific strength of autonomic modulation.

**Fig 1.**
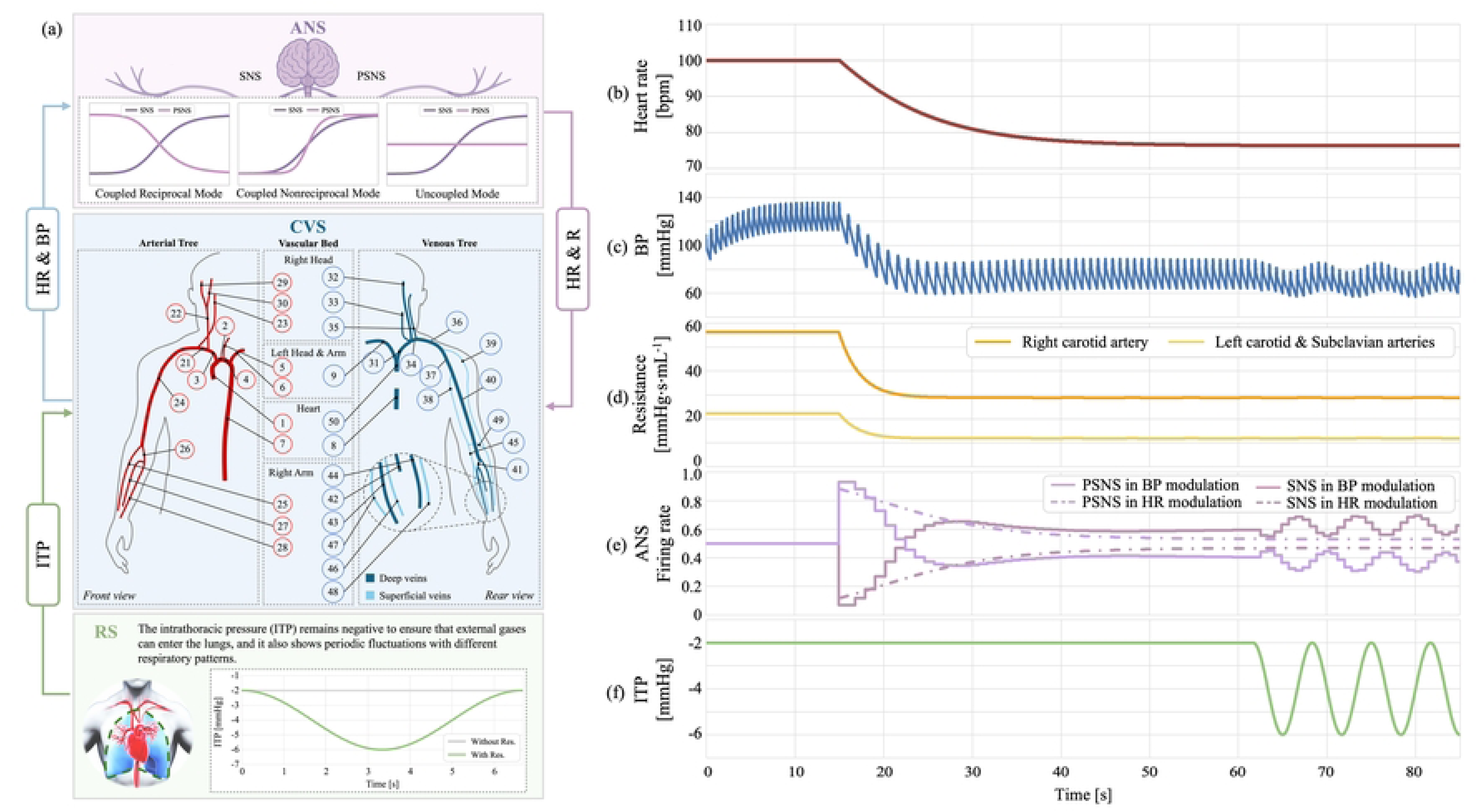
Computational model integrating the cardiovascular, respiratory, and autonomic nervous systems (CVS, RS, and ANS). (a) Schematic overview of the integrated structure of the CVS, RS, and ANS. HR and BP are transmitted to the ANS as fiber afferent activities, while HR itself and blood-flow resistance (R) are modulated by ANS regulatory signals. The R is the key determinant of BP dynamics in the model. ITP waves influence beat-to-beat BP dynamics over longer timescales. (b–f) Representative output signals generated by the model under closed-loop operation, shown across three intentionally designed phases to illustrate the functional contributions of the CVS, ANS, and RS modules rather than to represent distinct biologically occurring physiological stages. Phase I: Elevated initial levels of (b) HR, (c) BP, and (d) blood flow resistance, prescribed to demonstrate CVS-dominated dynamics under a high-load condition. Phase II: HR, BP, and blood flow resistance are gradually restored toward normal physiological ranges under (e) ANS regulatory signals that modulate SNS and PSNS activities, illustrating autonomic feedback regulation. Phase III: ITP oscillations generated by the RS module are introduced, contributing to cyclical variations observed in the BP trace and demonstrating respiration-driven modulation of hemodynamics after autonomic stabilization.

### 3.1 Cardiovascular system

The CVS module is responsible for reproducing HR and BP signals that vary in synchrony with the cardiac cycle. The CVS module was built on a closed-loop multiscale hemodynamic modeling framework that integrates hemodynamic model representing major arterial and venous structures, and lumped-parameter model representing cardiac chambers and peripheral arteriovenous interfaces [40],[41]. The governing equations are formulated as hemodynamic model approximations of the Navier–Stokes equations along the axial direction of the vessels. The pressure levels within vessels and heart chambers could be defined as:

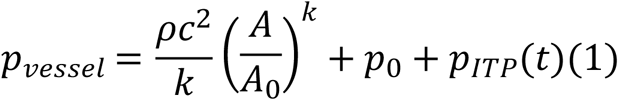

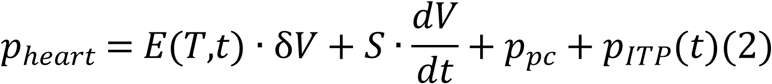

where *A* and *A*_0_ are the current and reference cross-sectional areas, respectively, and *k* is a stiffness exponent characterizing nonlinear wall elasticity. The parameter *c* is the Moens–Korteweg wave speed, reflecting pulse wave velocity under reference conditions, and *p*_0_ is the reference pressure. *ρ* is the blood density. *E* (*T*,*t*) · δ*V* represents the elastic component, *E*(*T*,*t*) is the time-varying elastance dependent on cardiac cycle *T* = 60/*HR* and time *t*, and δ*V* = *V* ― *V*_0_is the deviation from reference volume *V*_0_. The term *S* · *dV*/*dt* represents viscoelastic component, capturing the viscous dissipation caused by volume changes, where *S* is the myocardial viscoelastic coefficient, and *p*_*pc*_ denotes the pericardial pressure. The time-varying intrathoracic pressure, *p*_*ITP*_(*t*), is introduced to mimic the respiratory effects on pressure waveforms (see Section 3.2).

Vascular beds are represented using Kirchhoff’s circuit laws, with arterial and venous terminals treated as capacitive nodes connected by blood flow resistance. Volume dynamics are given by:

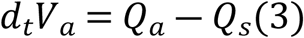

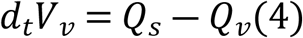

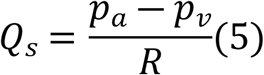

where *V*_*a*_and *V*_*v*_ are arterial and venous volumes, and *Q*_*a*_, *Q*_*v*_, and *Q*_*s*_are inflow from the upstream terminal arteries, the outflow toward the terminal veins, and exchange flow in vascular beds. The *p*_*a*_ and *p*_*v*_ are arterial and venous pressures. The *R* denotes the effective flow resistance of the vascular bed, regulating systemic BP levels.

The cardiovascular model combines chamber dynamics, vascular wall mechanics, and blood transport in the arterial and venous trees, enabling unified simulation of the time-varying pressure and flow. By specifying key physiological parameters such as cardiac cycle length, vessel geometry, and peripheral resistance, the model generates stable periodic cardiac dynamics, producing HR rhythms and synchronized BP oscillations. Simulated pressure waveforms in major vessels have been validated against physiological data, confirming that the model reproduces key hemodynamic characteristics of the human CVS.

### 3.2 Respiratory system

Under regular or controlled breathing conditions, respiration-related intrathoracic pressure (ITP) variations are primarily driven by the rhythmic contraction and relaxation of respiratory muscles. As a result, the temporal profile of ITP typically exhibits smooth and continuous low-frequency fluctuations dominated by a single principal frequency. Compared with the cardiac cycle, the respiratory cycle operates on a substantially longer timescale; therefore, its influence on cardiac chamber pressures and vascular pressures can be regarded as a slowly varying external modulatory component. Based on these physiological characteristics, and in line with previous related work [40],[41], the temporal variation of ITP, *p*_*ITP*_(*t*), is approximated using a simple periodic function:

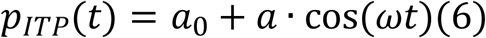

where *a*_0_denotes the baseline respiration-related BP wave, and the angular frequency *ω* and amplitude *a* are specified by the selected respiratory pattern. This term is further incorporated into the analytical formulation of BP to characterize respiration-induced BP fluctuations.

### 3.3 Autonomic nervous system

#### 3.3.1 Autonomic modulation modeling

The HR signals generated by the CVS module are transmitted to the central nervous system as HR-related afferent fiber activations. BP regulation is modeled based on baroreceptor-mediated mechanism. Three baroreceptors, located in the aortic arch and bilateral carotid sinuses, detect pressure fluctuations in the aorta (*P*_*a*_), left and right carotid arteries (*P*_*l*_ & *P*_*r*_), corresponding to arteries No. 2, 5, and 22 in the CVS module (Fig 1). The baroreceptors are assumed to have identical physiological properties in pressure sensing and signal transduction. Accordingly, the average real-time pressure across the aorta and bilateral carotid arteries is computed and used as the afferent input signal for BP regulation.

Modulation of SNS and PSNS activities was implemented using sigmoid functions to capture their nonlinear responsiveness to HR and BP states in parallel pathways. For each pathway *i* ∈ {*HR*, *BP*}, the activation levels are defined as:

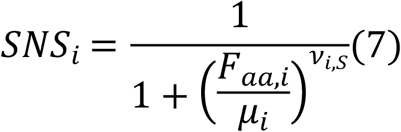

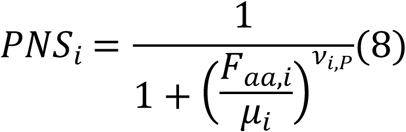

where *F*_*aa*,*HR*_ = *HR*/60 represents the current HR in beats per second (bps), and *F*_*aa*,*BP*_ = (*P*_*a*_ + *P*_*l*_ + *P*_*r*_)/3 represents the average BP across the aortic arch and carotid sinuses. The parameters *v*_*i*,*S*_ and *v*_*i*,*P*_ control the response slopes of the sympathetic and parasympathetic branches, respectively, and *μ*_*i*_ denotes the baseline HR or BP levels representing the target level. The parameter *μ*_*HR*_was determined as the mean of the measured human data, while *μ*_*BP*_was derived from MAP. MAP was used as the primary index for evaluating BP regulation. For each simulation, systolic blood pressure (SBP) and diastolic blood pressure (DBP) were extracted from the model-output pressure waveforms, and MAP was computed as:

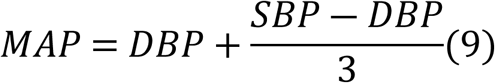

The ANS modulation signal was computed as a weighted combination of *SNS*_*i*_ and *PNS*_*i*_:

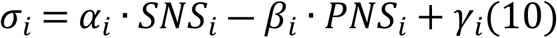

where *α*_*i*_ and *β*_*i*_represent the relative activity levels of SNS and PSNS, respectively, and *γ*_*i*_is the basal activation level of target organs in the absence of ANS modulation. The target parameters for the control loops were guided by the primary determinants of HR and BP in the CVS submodule. For HR modulation, the target parameter is the HR itself (bps), while for BP modulation, the target parameter is the total peripheral blood flow resistance (R) of the arterial tree. The modulatory dynamics of each target parameter were governed by a first-order ordinary differential process as:

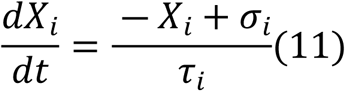

where *X*_*i*_ denotes the target parameter: HR itself for HR modulation (*X*_*HR*_ = *HR*) and R for BP modulation (*X*_*BP*_ = *R*), with *τ*_*i*_ being the characteristic time constant (s). For HR modulation, *τ*_*HR*_ = 25s, and for BP modulation, *τ*_*BP*_=5 s. In addition to direct ANS modulation, BP modulation was also influenced by HR-driven changes in cardiac output and by respiratory effects, yielding a physiologically integrated model.

#### 3.3.2 Autonomic nervous system functions

Building on the autonomic regulation framework, this study characterized ANS function by separately parameterizing autonomic control modes and the relative activity of the SNS and PSNS. Autonomic control modes define the structural patterns of SNS–PSNS coordination that govern cardiovascular regulation, whereas relative activity refers to the branch-specific scaling of sympathetic and parasympathetic outputs specified at the beginning of each simulation. This formulation allows the model to distinguish between qualitative differences in autonomic regulation strategies and quantitative differences in the strength of SNS and PSNS modulation acting on different interoceptive signals.

Specifically, autonomic control modes were first systemically defined within the ANS submodule to represent distinct SNS–PSNS coordination schemes governing HR and BP regulation under different external stimuli and physio/psychological conditions. Three types of autonomic control modes were defined: coupled reciprocal, coupled nonreciprocal, and uncoupled. Fig 2 illustrates how ANS activity patterns vary with different levels of afferent fiber activation (*F*_*aa*,*i*_) under each control mode. The coupled reciprocal mode [Fig 2(a)], one of the earliest forms of autonomic control described in the previous literature, is characterized by an opposing activity between the SNS and PSNS branches, such that activation of one division is accompanied by inhibition of the other. In the model, reciprocal modes were represented by assigning opposite signs to the slope parameters *v*_*i*_ of SNS and PSNS. Following the parameter settings in [43], the response slopes of SNS and PSNS were set to ± 7 to produce responses of equal magnitude and opposite direction. In coupled nonreciprocal mode [Fig 2(b)], the *v*_*i*_ values of SNS and PSNS were assigned the same sign: positive slopes indicate coinhibition, whereas negative slopes indicate coactivation. In the uncoupled mode [Fig 2(c)], one division was modeled by setting its *v*_*i*_value to zero, effectively withdrawing its response so that its activity remained constant across all *F*_*aa*,*i*_ levels. In addition to specifying control modes, the model allowed independent adjustment of the relative levels of SNS and PSNS activity using weighting parameters. Meanwhile, within each predefined control mode, the relative activity levels of the SNS and PSNS branches could be independently adjusted at the beginning of model-based simulations, thereby determining the effective gain of sympathetic or parasympathetic modulation on HR and BP without altering the underlying control structure. The weighting factors *α*_*i*_ and *β*_*i*_ scaled the respective contributions of SNS and PSNS outputs to modulation of each target parameter. These parameters controlled the relative gain of sympathetic and parasympathetic influences while preserving the predefined SNS–PSNS interaction pattern associated with each control mode. To examine how variations in autonomic balance influenced HR and BP dynamics, *α*_*i*_ and *β*_*i*_were varied from 0 to 1 in increments of 0.2. As shown in Fig 2, the relative activity levels of SNS and PSNS within each control mode were modulated by tuning *α*_*i*_and *β*_*i*_ within this normalized range.

**Fig 2.**
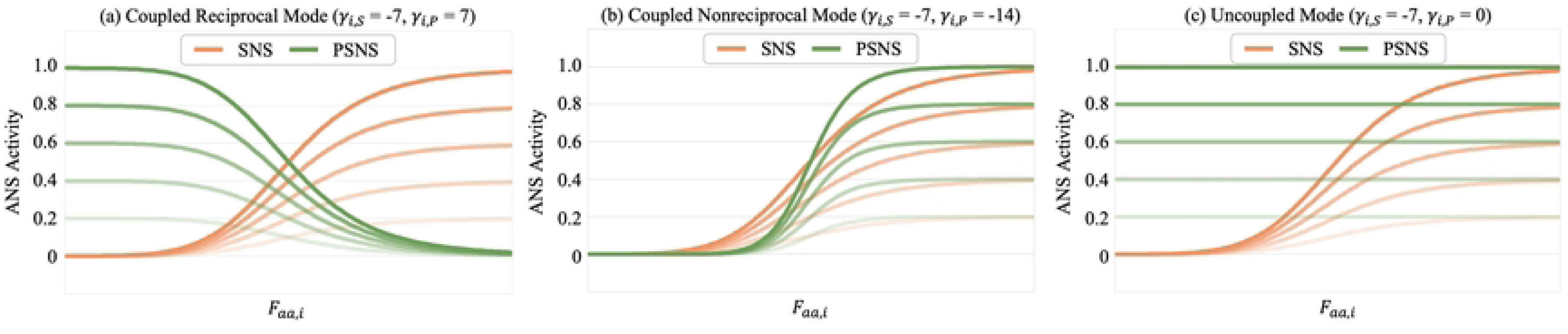
Definition of autonomic control modes and corresponding SNS–PSNS activity patterns as a function of afferent fiber activation. The model defines three autonomic control modes based on the relative interaction between SNS and PSNS branches. (a) Coupled reciprocal mode, in which SNS and PSNS exhibit opposing activation patterns with response slopes of equal magnitude but opposite sign. (b) Coupled nonreciprocal mode, in which SNS and PSNS share the same response sign, resulting in coactivation or coinhibition depending on slope direction. (c) Uncoupled mode, in which the response of one autonomic branch is withdrawn by setting its slope to zero, rendering its activity invariant across afferent input levels. Within each mode, the relative activity levels of SNS and PSNS are further modulated by the weighting parameters *α*_*i*_ and *β*_*i*_, which scale their respective contributions to autonomic regulation.

### 3.4 Output waveforms

Fig 1(b)–(f) illustrate representative waveforms generated by the model under closed-loop operation, including HR, BP, R, ANS outputs, and ITP. To clearly demonstrate the functional roles of the CVS, ANS, and RS modules within the integrated framework, the simulation timeline was intentionally divided into three designed stages, each highlighting the dominant contribution of a specific regulatory subsystem. These stages were defined for methodological illustration purposes rather than to represent distinct biologically occurring physiological states. The simulation was therefore structured to systematically characterize the dynamic responses of the CVS, RS, and ANS across these predefined regulatory phases. Accordingly, the time course was divided into three functionally motivated stages: (i) an initialization stage dominated by cardiovascular load, (ii) a regulation stage driven by autonomic feedback, and (iii) a modulation stage dominated by respiratory rhythms.

In Phase I (0–15 s), the model was initialized with elevated HR, BP, and R to simulate a high-load physiological state following external stimuli or internal stress. HR was set to a high level (100 bpm), and R remained elevated, reflecting significant vasoconstriction in peripheral vessels. The increases in HR and R jointly raised BP, which also exhibited cardiac cycle-synchronized oscillations. During Phase I, the ANS was inactive and no respiratory-induced ITP fluctuations were applied. This stage was designed to isolate the CVS response under a prescribed high-load condition, providing a reference state for the subsequent initiation of autonomic regulation.

In Phase II (15–65 s), the elevated HR and BP were transmitted as afferent inputs to the ANS module, initiating autonomic feedback regulation. SNS output gradually decreased, whereas PSNS output increased. Consequently, HR and R progressively declined, leading to recovery of BP toward the normal physiological range. During this process, SNS and PSNS outputs approached equilibrium, characterizing a typical coupled reciprocal pattern of ANS regulation. This phase demonstrates the role of the ANS module in restoring cardiovascular variables toward stable operating ranges through feedback control.

In Phase III (65–85 s), periodic ITP fluctuations generated by the RS module were introduced, simulating respiratory modulation of cardiac output. The rhythmic negative ITP waves induced low-frequency respiratory modulation in the BP waveform, superimposed on the higher-frequency cardiac oscillations. After ANS-mediated recovery, the mean BP level further decreased, and the ANS outputs exhibited mild fluctuations synchronized with the respiratory rhythm. This stage explicitly illustrates how respiratory dynamics, introduced after autonomic stabilization, modulate hemodynamic signals on a slower timescale.

Together, the three designed phases illustrate the dynamic behavior of the computational model across multiple regulatory timescales, with each phase emphasizing the contribution of a specific subsystem rather than depicting sequential biological stages. These simulations demonstrate the capacity of the closed-loop integration of CVS, RS, and ANS modules to reproduce key temporal characteristics of interoceptive dynamics.

## 4. Experiment 1: HR and BP responses in TD and ASD under external stimulation

### 4.1 Experimental setting

External stimulation induces rapid shifts in ANS function, reflected in changes in autonomic control modes and relative SNS–PSNS activity. To evaluate autonomic regulatory capacity, HR and BP responses were simulated across different control modes and SNS–PSNS activity combinations using the computational model. The simulated responses were compared with representative empirical ranges of HR and BP reported in previous studies of TD and ASD populations, enabling identification of control modes that reproduce characteristic physiological responses under stimulation.

Previously reported HR and BP responses observed during the head-up tilt (HUT) test [45] were selected as the empirical basis for this study and used to guide model configuration. The HUT test was chosen because it provides a standardized experimental paradigm for assessing autonomic regulation under gravitational perturbation in clinical research. As a passive postural challenge, HUT minimizes confounding effects from voluntary movement, skeletal muscle activation, and cognitive or emotional strategies. The test typically involves a rapid transition from a supine to an upright position (≈60–70°), inducing characteristic SNS– PSNS adjustments that support HR elevation and BP stabilization in response to postural challenge.

Real measured HR and BP levels before and after the HUT test (Table 1) were used to parameterize afferent fiber activations (*F*_*aa*,*i*_) and baseline target levels (*μ*_*i*_) of HR and BP, respectively, for the ANS modulatory loop in the sigmoid-based framework. Median measured values at supine rest in TD and ASD groups were used to define afferent input variables (*F*_*aa*,*i*_) serving as the initial inputs to the model. In contrast, uniform baseline values derived from the median post-HUT measurements of the TD group were applied to both groups, providing a common reference target and enabling direct comparison of autonomic regulation between TD and ASD individuals. Full parameter settings are summarized in Table 2.

**Table 1.** Real measured heart rate (HR) and systolic and diastolic blood pressure (SBP & DBP) ranges at supine rest and HUT test conditions in TD and ASD groups.

| Groups | Interoceptive signals | Supine Rest | HUT Test |
| --- | --- | --- | --- |
| TD | HR [bpm] | 54.51–71.49 | 69.01–86.99 |
|  | SBP [mmHg] | 109.51–134.19 | 99.49–132.51 |
|  | DBP [mmHg] | 60.16–71.84 | 58.69–79.31 |
|  | MAP [mmHg] | 76.61–92.62 | 72.29–97.04 |
| ASD | HR [bpm] | 62.34–91.66 | 82.54–115.46 |
|  | SBP [mmHg] | 106.98–129.02 | 105.5–130.5 |
|  | DBP [mmHg] | 60.63–77.37 | 65.89–82.11 |
|  | MAP* [mmHg] | 76.08–94.59 | 79.09–98.24 |

\* MAP: mean arterial pressure calculated via Eq. (9).

**Table 2.** Baseline targets (*μ*) and afferent activations (*F*_*aa*_) for HR and BP in TD and ASD groups.

| Group | $\mu_{HR}$ [bps] | $\mu_{BP}$ [mmHg] | $F_{aa,HR}$ [bps] | $F_{aa,BP}$ [mmHg] |
| --- | --- | --- | --- | --- |
| TD | 1.300 | 116 | 1.050 | 84.66 |
| ASD | 1.300 | 116 | 1.283 | 85.33 |

### 4.2 Results

#### 4.2.1 Autonomic regulation in TD and ASD groups

Three-dimensional (3D) surface plots were used to illustrate HR and BP responses in TD and ASD groups and to demonstrate how computational modeling, combined with empirical measurements, clarifies ANS regulatory characteristics. Fig 3 illustrates HR and BP responses in TD and ASD groups under three autonomic control modes during HUT stimulation. A set of representative ANS activity curves shown in Fig 3 (a) was generated using maximal weighting parameters for both autonomic branches, with *α*_*i*_ and *β*_*i*_ fixed at 1.0. Relative SNS and PSNS activity levels were systematically varied from 0 to 1.0 in increments of 0.2 to construct the response surfaces. The 3D plots in Fig 3 (b)-(j) display simulated steady-state HR, SBP and DBP responses across SNS–PSNS relative activity weights, with blue circular markers indicating TD group outcomes and red square markers indicating ASD group outcomes following HUT stimulation. In addition, the blue and red cross markers on the simulated 3D surfaces indicate HR and BP responses falling within the empirically observed ranges of TD and ASD groups, while the corresponding color-coded polygons delineate the SNS–PSNS relative activity regions in which simulated responses aligned with experimental measurements.

**Fig 3.**
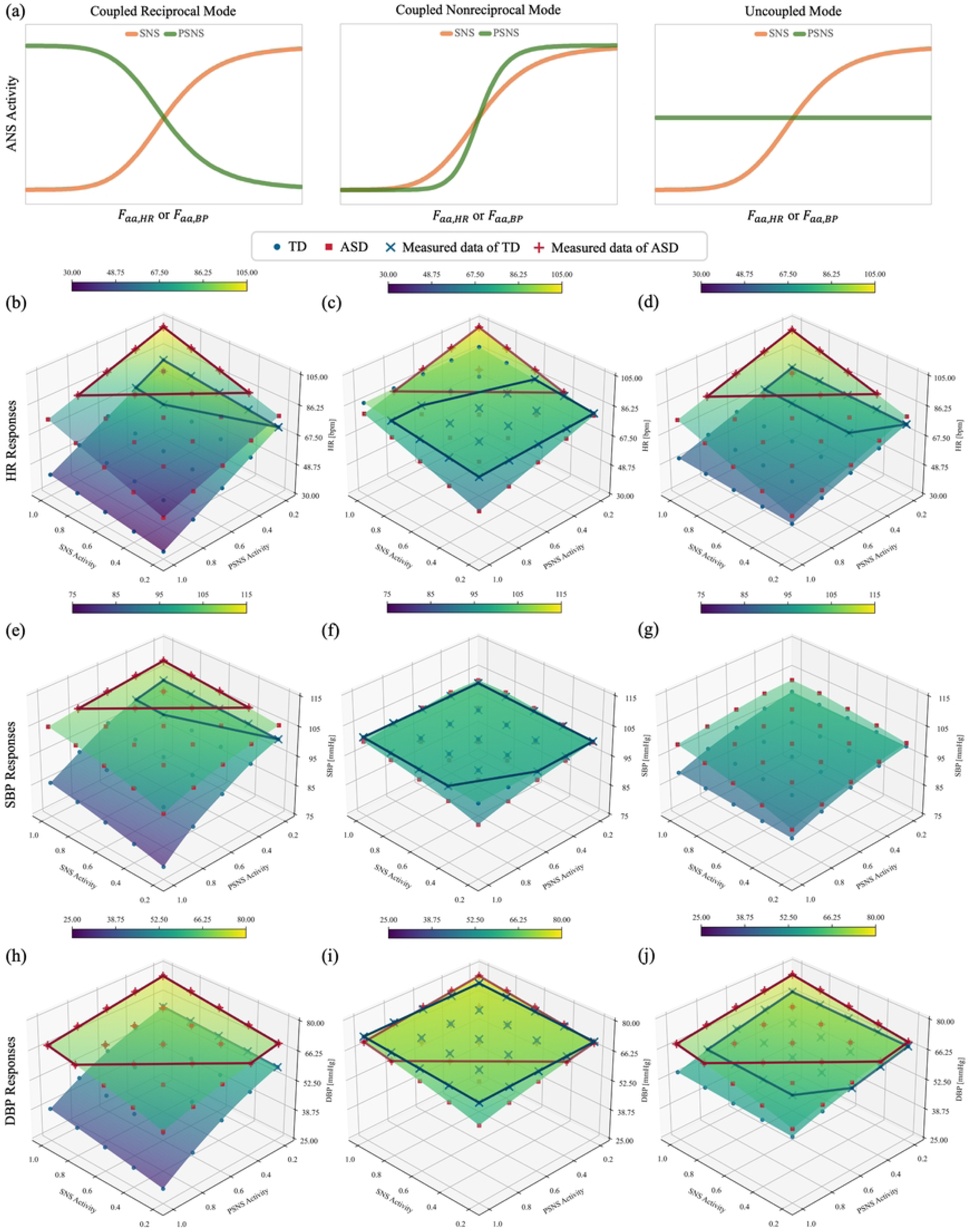
Simulated HR and BP responses of TD and ASD groups across three autonomic control modes during HUT stimulation. The upper panels illustrate the autonomic response profiles for the coupled reciprocal, coupled nonreciprocal, and uncoupled modes, generated with maximal weighting parameters for both SNS and PSNS (*α*_*i*_ = *β*_*i*_ = 1.0). The three rows of 3D blue and red dotted surfaces present simulated steady-state HR, SBP, and DBP responses of TD and ASD groups, respectively, across the SNS–PSNS activity plane. Blue markers denote simulated TD responses that fall within the empirically measured HR and BP ranges, and red markers denote the corresponding ASD responses. The spatial distributions of markers across the surfaces highlight the combinations of SNS and PSNS relative activities for which the simulated outcomes align with empirical observations. Across the three autonomic control modes, ASD individuals exhibited higher HR, SBP, and DBP levels and greater sensitivity to changes in SNS activity compared with TD individuals, whereas TD responses aligned more closely with expected physiological patterns under the coupled reciprocal mode. In contrast, responses from the ASD group were distributed in similar SNS–PSNS regions across all three modes, indicating reduced differentiation in their autonomic regulatory behavior.

Under both the coupled reciprocal mode [Fig 3(b)] and the uncoupled mode [Fig 3(d)], empirical HR data from both TD and ASD groups were distributed within regions characterized by high SNS weighting and low PSNS weighting. Compared with the TD group, the ASD group exhibited a greater concentration of data points in the high SNS region, as well as additional distribution extending toward regions with relatively higher PSNS weighting, indicating that the simulated HR responses corresponding to ASD observations were associated with comparatively higher relative PSNS activity. In the coupled nonreciprocal mode [Fig 3(c)], empirical HR data from the ASD group remained clustered in regions with high SNS and low PSNS weighting, whereas TD group data shifted toward regions with lower SNS and higher PSNS weighting.

The distribution of empirical BP responses was analyzed in a manner analogous to HR, as illustrated in Fig 3(e)–(j). Because SBP and DBP jointly reflect blood pressure regulation, the overlap between their respective SNS–PSNS activity regions was taken to indicate the relative SNS–PSNS activity combinations capable of modulating BP levels. In the coupled reciprocal mode [Fig 3(e) and (h)], the overlap between the SBP and DBP regions collapses to a narrow line corresponding to a PSNS weight of 0.2, which characterizes the SNS–PSNS relative activity region of the TD group. In contrast, for the ASD group, the overlap region coincides with the red polygon identified in Fig 3(e), indicating the SNS–PSNS relative activity region associated with BP regulation. In the coupled nonreciprocal mode [Fig 3(f) and (i)], the SNS–PSNS relative activity region associated with BP regulation could be identified only from the empirical DBP region for the ASD group. In the uncoupled mode [Fig 3(g) and (j)], the SNS–PSNS relative activity regions for both TD and ASD groups were identifiable solely based on the empirical DBP regions. Comparison of BP-based analyses indicates that, in the coupled reciprocal mode, the overlap SNS–PSNS relative activity region of the TD group is concentrated in areas of high SNS and low PSNS weighting, whereas the corresponding region for the ASD group shows an extension toward higher PSNS weighting. This pattern is consistent with that observed in the HR-based analysis. In contrast, under the coupled nonreciprocal and uncoupled modes, the SNS–PSNS relative activity regions of both TD and ASD groups are broader.

Notably, using HR as an example, the relatively higher PSNS activity indicated by the model for the ASD group appears inconsistent with the overall higher post-HUT HR levels observed empirically. The ASD group exhibited higher empirically measured resting HR, which was used as the initial afferent input to the model. This elevated baseline resulted in higher SNS and PSNS activation values at the onset of modulation and an upward shift of the simulated HR response surfaces. Consequently, steady-state HR after HUT remained higher in the ASD group across autonomic weighting parameters. In this study, SNS and PSNS activation values reflect instantaneous neural responses to afferent cardiovascular signals, whereas the weighting parameters α and β quantify the relative effectiveness of autonomic output. Accordingly, the elevated post-HUT HR in the ASD group is attributable to higher initial cardiovascular states, rather than increased sympathetic dominance or parasympathetic suppression.

#### 4.2.2 Direction and magnitude of ANS regulation

The general trajectories of interoceptive responses following the HUT test were evaluated under four extreme combinations of SNS and PSNS weighting parameters (*α* and *β*) to characterize the roles of these parameters in determining the direction and magnitude of ANS-modulated interoceptive regulation. As shown in Fig 4(a), ANS-modulated HR outcomes, for example, varied markedly across weighting conditions depending on the dominant autonomic branch in TD individuals. When PSNS weighting was minimal (*β* = 0.2), HR exhibited a upward regulatory direction. In contrast, when PSNS weighting was dominant (*β* = 1.0), HR showed a continuous decline toward the lower level than baseline. However, according to Eq. (11), the direction and magnitude of HR adjustments in the model were governed directly by the modulatory signal (*σ*), rather than by the weighting parameters themselves. Fig 4(b)–(e) present the temporal evolution of both the HR afferent activation level [*F*_*aa*,*HR*_; 1.05 bps (63 bpm), in Table 2] and *σ* under the four weighting conditions. In the configurations shown in Fig 4(b) and (c), *F*_*aa*,*HR*_exceeded *σ* at the onset of autonomic engagement. Under these conditions, the lower *σ* level imposed a downward regulatory influence, driving both *F*_*aa*,*HR*_and *σ* to decrease concurrently until the difference between the two signals was eliminated. HR stabilization was achieved once *F*_*aa*,*HR*_ and *σ* converged, resulting in a reduced post-modulation HR level. In contrast, in the configurations shown in Fig 4(d) and (e), *σ* was initially higher than *F*_*aa*,*HR*_. The elevated *σ* level generated a sustained upward regulatory drive, leading to progressive HR elevation until *σ* and *F*_*aa*,*HR*_ approached equivalence, and stabilization was attained. Across the weighting conditions examined in Fig 4(b)–(e), the direction of HR variation was determined by the sign of the difference between *σ* and *F*_*aa*,*HR*_, whereas the magnitude of HR elevation or reduction scaled with the absolute value of this difference. In these simulations, SNS and PSNS activities influenced HR responses by modulating the *σ* signal, rather than acting as independent drivers of HR regulation.

**Fig 4.**
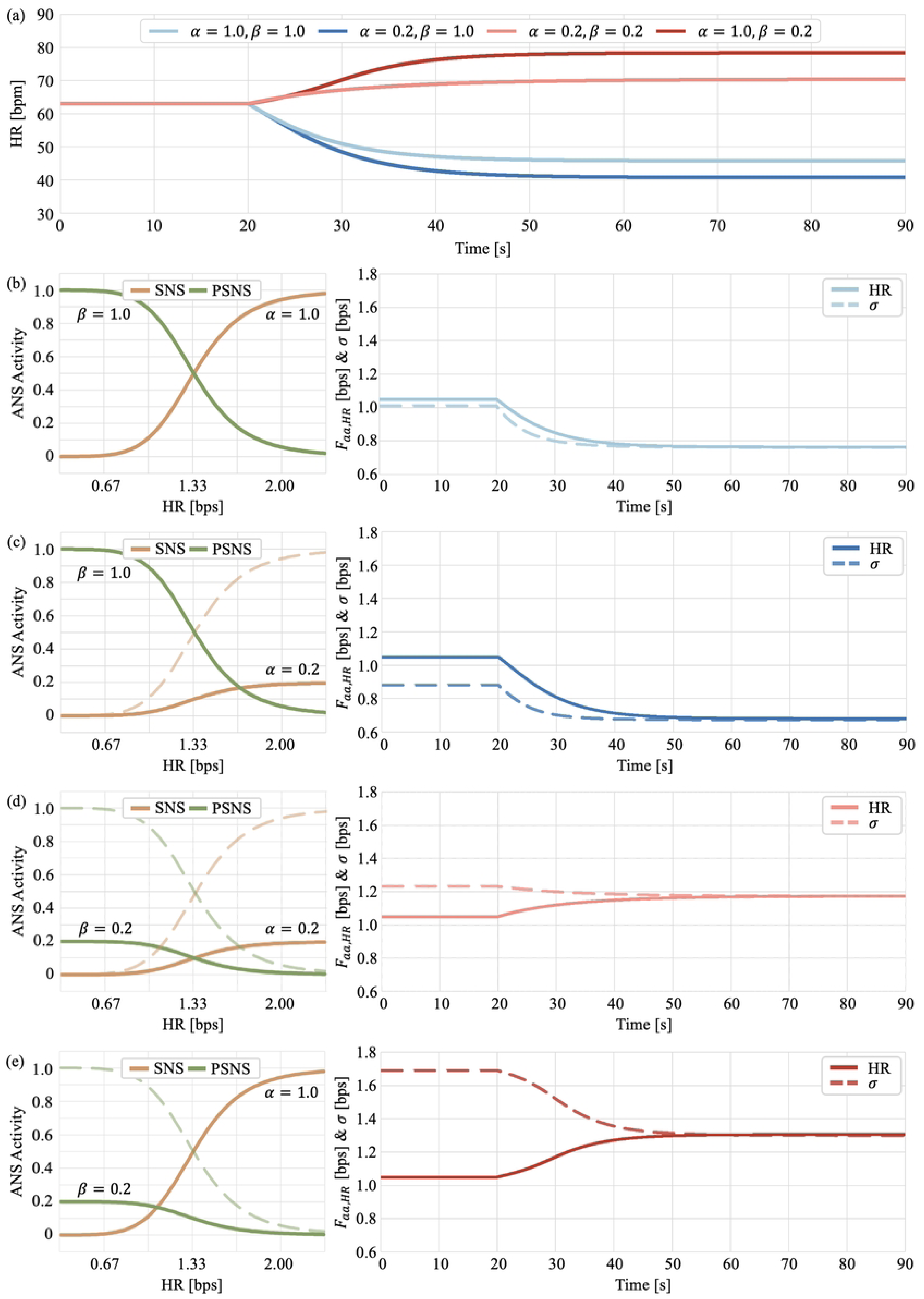
Time-dependent HR responses and underlying autonomic modulation mechanisms following the HUT test under four extreme SNS–PSNS weighting combinations (α, β) in TD individuals. (a) Simulated HR trajectories during ANS modulation for different combinations of SNS and PSNS weights. (b)–(e) Mechanistic analyses illustrating the temporal evolution of the afferent activation level of HR (*F*_*aa*,*HR*_) and the modulation signal *σ* under the corresponding weighting conditions shown in (a). When *F*_*aa*,*HR*_ exceeds σ at the onset of autonomic engagement (b, c), the lower *σ* level imposes a downward regulatory drive, causing both *F*_*aa*,*HR*_ and *σ* to decrease concurrently until convergence at a lower stabilized HR level. In contrast, when *σ* is initially higher than *F*_*aa*,*HR*_ (d, e), the elevated *σ* level generates a sustained upward regulatory drive, leading to progressive HR elevation until convergence at a higher stabilized level.

#### 4.2.3 Baseline definition for TD and ASD groups

This study further examined the consequences of defining group-specific baseline values based on empirically measured post-HUT HR or BP levels, in comparison with using unified baseline values across TD and ASD groups. Fig 5 illustrate the baseline HR selection under coupled reciprocal mode and simulated HR responses. The unified baseline *μ*_*HR*_ = 1.300 bps (78 bpm) effectively reveals marked differences in the initial regulatory state [Fig 5(a)] within the SNS and PSNS activation levels [Eqs. (7) and (8)]. The TD group exhibited an F_*aa*,*HR*_ of 1.050 bps (63 bpm), which was associated with relatively low SNS activity and high PSNS activity (SNS = 0.183, PSNS = 0.817). In contrast, the ASD group exhibited an F_*aa*,*HR*_of 1.283 bps (77 bpm), and under the same baseline, showed substantially elevated SNS activity and reduced PSNS activity (SNS = 0.477, PSNS = 0.523), reflecting a characteristic pattern of autonomic imbalance. Assigning the ASD post-HUT HR value of 1.650 bps (99 bpm) as a group-specific baseline would lead the model to interpret this elevated HR as the intended regulatory target. Under this condition, the resulting SNS and PSNS estimates (SNS = 0.147, PSNS = 0.853) would fall within an apparently typical range, thereby obscuring abnormalities in ASD autonomic regulation. When *μ*_*HR*_ is set to 99 bpm, the simulated ASD HR response under the coupled reciprocal mode becomes lower than that of the TD group, and no empirical ASD HR values fall within the corresponding response surface. Elevated baseline values reduce initial sympathetic drive and prematurely enhance parasympathetic activation, leading to a markedly restricted regulatory range. Baseline values anchored to abnormally elevated physiological states therefore lack biological validity and distort simulated regulatory trajectories by artificially normalizing characteristic autonomic features in ASD. Such baseline adjustment would further compromise comparability between groups by eliminating a shared regulatory target, making it impossible to evaluate regulatory deviation or efficiency across conditions.

**Fig 5.**
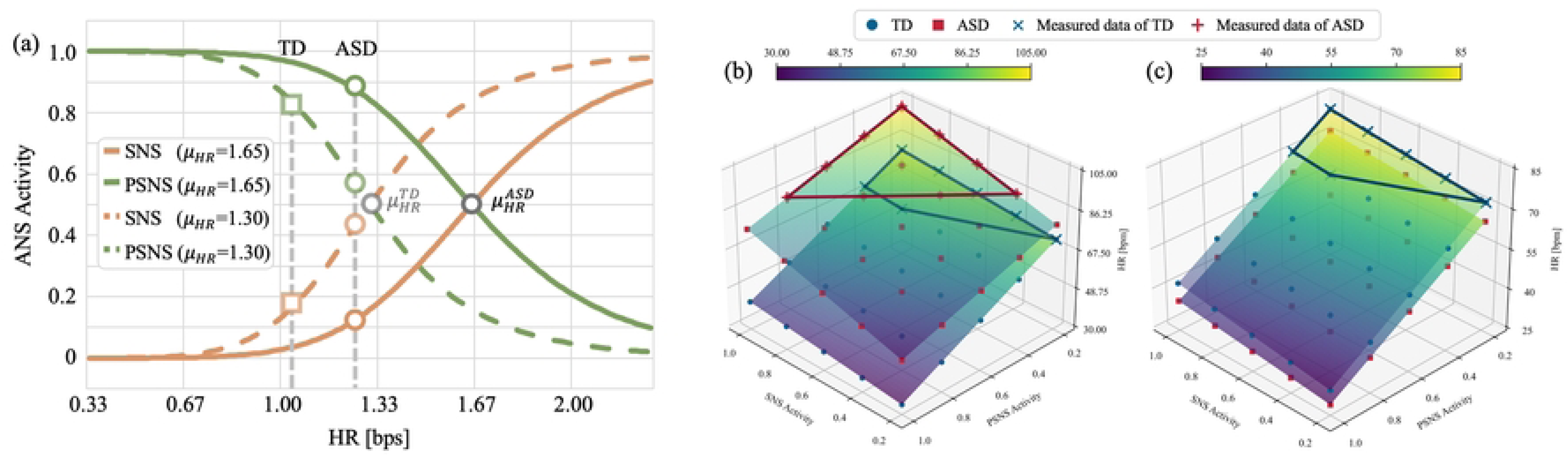
Effects of baseline selection on autonomic activity estimates and simulated HR responses. (a) Sympathetic (SNS) and parasympathetic (PSNS) activities derived from the sigmoid-based autonomic modulation function under two baseline settings: *μ*_*HR*_= 1.30 bps (78 bpm; solid lines) and *μ*_*HR*_ = 1.65 bps (99 bpm; dashed lines). The TD resting HR (63 bpm) and ASD resting HR (77 bpm) produce distinct Fₐₐ values, which yield clearly differentiated SNS–PSNS balances when a unified baseline is applied. Assigning the ASD post-HUT HR (99 bpm) as a group-specific baseline shifts the predicted autonomic state toward a falsely normalized pattern and masks characteristic ASD autonomic imbalance. (b) Simulated HR surfaces across SNS–PSNS weighting combinations under the unified baseline (*μ*_*HR*_= 1.30 bps), overlaid with measured HR values from TD and ASD individuals following the HUT test. The TD data cluster aligns with the physiologically expected high-SNS/low-PSNS region, whereas ASD data extend toward elevated PSNS relative activity. (c) Simulated HR surfaces generated using the ASD-specific elevated baseline (*μ*_*HR*_= 1.65 bps). The increased baseline compresses the effective regulatory range, resulting in attenuated HR responses that fail to capture empirical ASD data distribution patterns, thereby illustrating how an abnormally high baseline distorts simulated autonomic trajectories.

Overall, TD individuals exhibited clearly differentiated SNS–PSNS relative activity patterns across autonomic control modes, enabling identification of a dominant regulatory strategy recruited in response to external postural challenge. In contrast, ASD individuals showed greater similarity in SNS–PSNS activity distributions across the three control modes, suggesting reduced differentiation in autonomic control. This convergence implies that, when exposed to the same external stimulus, ASD individuals may not consistently engage a stable or mode-specific ANS regulatory strategy to modulate interoceptive signals. Within this context, the coupled reciprocal mode demonstrated greater potential to capture intrinsic autonomic regulatory processes in TD individuals. Under this mode, empirical HR and BP values in the TD group were primarily distributed within regions characterized by high SNS and low PSNS relative activity, which is consistent with the expected physiological response to HUT stimulation. In contrast, although most ASD observations were likewise located within regions of elevated SNS activity, their distributions extended toward areas with relatively higher PSNS weighting. This shift suggests that PSNS activity in ASD individuals may not be effectively suppressed following HUT stimulation. Such persistent parasympathetic engagement may constrain ANS-mediated interoceptive adjustment, thereby reducing the adaptability of cardiovascular regulation to external perturbations.

## 5. Experiment 2: Respiratory effects on ANS-modulated BP restoration

### 5.1 Experimental setting

The present study also investigated the modulatory impact of respiration on ANS-regulated BP restoration process. The model was initialized at a relatively high BP level, reflecting a physiological condition in which blood pressure has already been elevated by internal variability or prior external challenges. The MAP level [Eq. (9)] were selected as key metrics and systematically examined under varying combinations of SNS– PSNS relative activity weights. Physiologically, once the internal or external challenges are relieved, elevated BP is typically reduced through baroreflex-mediated autonomic regulation. This recovery process is known to involve a coupled reciprocal mode [17],[19], facilitating the gradual restoration of BP toward normal physiological levels. Accordingly, respiratory effects on BP restoration were examined under the coupled reciprocal mode, in which sympathetic activity is progressively suppressed while parasympathetic activity is concurrently enhanced.

The model was further examined under three respiratory conditions during BP restoration [Fig 6]: (1) without respiratory influence; (2) under a normal respiratory pattern, characterized by relatively low amplitude and high frequency (15 bpm); and (3) under a deep respiratory pattern, characterized by higher amplitude and lower frequency (9 bpm). Under this setting, respiratory-related ITP fluctuations [Eq. (6)] followed the patterns summarized in Table 3.

**Fig 6.**
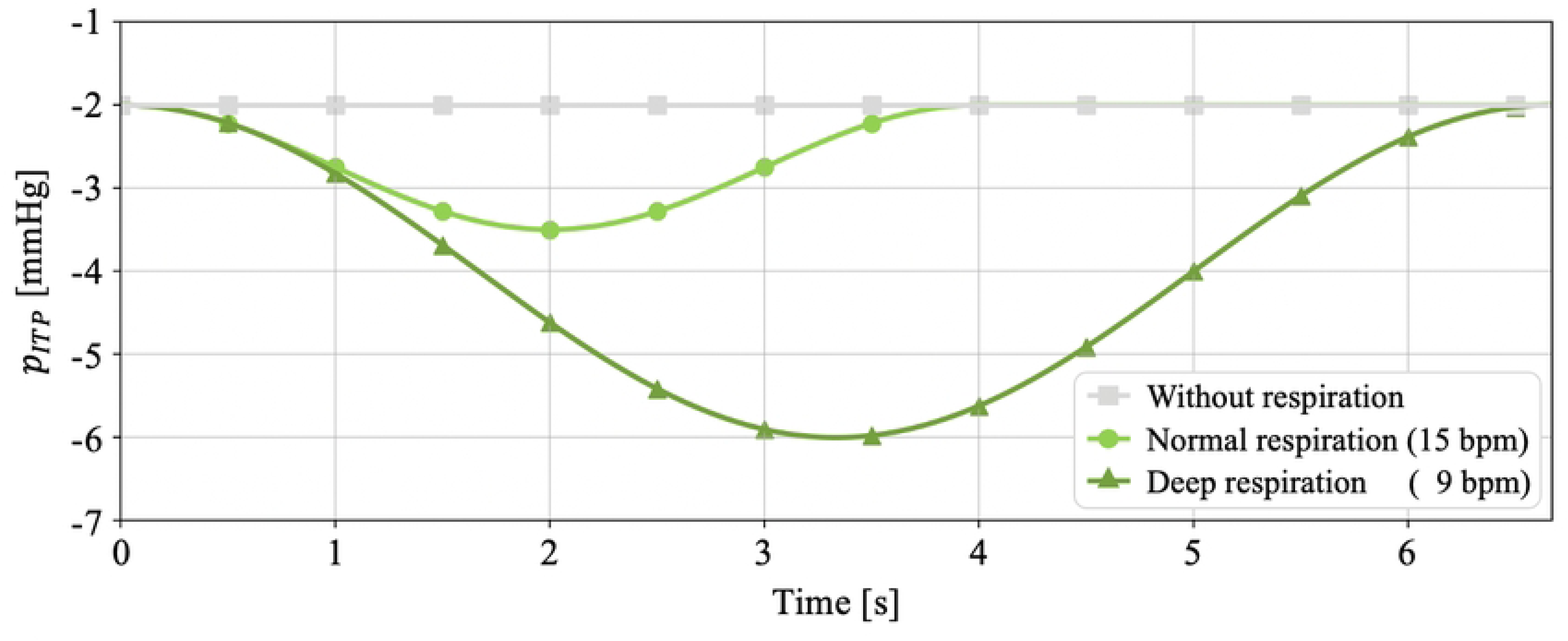
Single-cycle intrathoracic pressure (ITP) waveforms under three respiratory conditions during BP restoration. Respiratory-related ITP fluctuations are shown for (1) without respiration, (2) normal respiration characterized by low amplitude and high frequency (15 bpm), and (3) deep respiration characterized by higher amplitude and lower frequency (9 bpm).

**Table 3.** Parameter settings for ITP fluctuations, *p*_*ITP*_(*t*), under different respiratory conditions.

| Conditions | $a_0$ [mmHg] | $a$ [mmHg] | $\omega$ [rad/s] |
| --- | --- | --- | --- |
| Normal respiration | -2.75 | 0.75 | 1.57 [15 bpm] |
| Deep respiration | -3.00 | 2.00 | 0.94 [9 bpm] |
| Without respiration * | -2.00 | 0 | 0 |

\* For the “Without respiration”, a constant baseline offset (-2.00 mmHg) was retained to represent a static ITP level in the absence of respiratory oscillations.

### 5.2 Results

Fig 7 shows the stabilized MAP levels after ANS modulation across combinations of SNS and PSNS relative activities, with *α* and *β* ranging from 0 to 1. Under normal respiration [Fig 7(b)], the final MAP level was comparable to that observed in the absence of respiratory influence [Fig 7(a)], whereas deep breathing resulted in an overall lower MAP level [Fig 7(c)]. For instance, when both SNS and PSNS activities were set to 0.6, MAP values were similar between the no-respiration (101.68 mmHg) and normal breathing (100.10 mmHg) conditions, whereas a marked reduction was observed under deep breathing (94.58 mmHg). This pattern indicates that respiratory activity can further increase the magnitude of BP reduction during ANS-mediated restoration, with larger ITP amplitudes and longer oscillation periods, such as those observed during deep respiration, being associated with lower final MAP levels.

**Fig 7.**
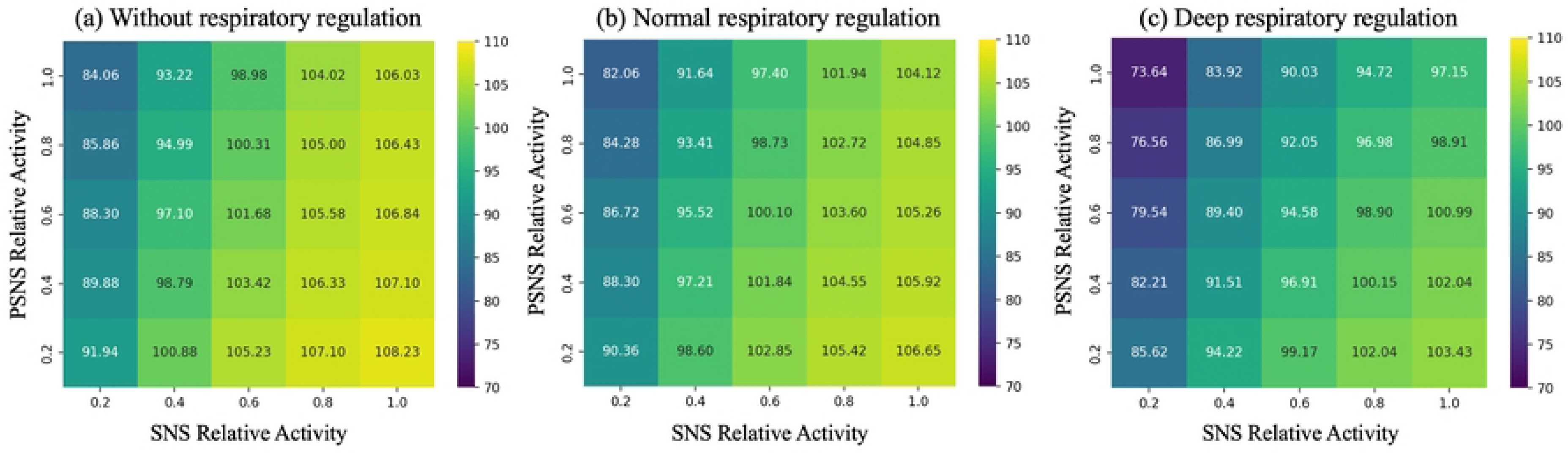
Respiratory modulation of ANS-mediated MAP restoration across SNS–PSNS activity space. Stabilized MAP following ANS modulation across combinations of relative SNS and PSNS activities (0–1) under (a) absence of respiratory modulation, (b) normal respiration, and (c) deep respiration. Normal respiration produced MAP levels comparable to those without respiratory influence, whereas deep respiration resulted in an overall downward shift in MAP. Under conditions of high SNS activity, increasing PSNS weighting was associated with a more pronounced MAP reduction under deep respiration than under the other respiratory conditions.

Under SNS hyperactivity (e.g., SNS = 1.0), distinct MAP regulation patterns emerged depending on respiratory modulation. As PSNS relative activity increased from 0.2 to 1.0, a clear decreasing trend in MAP was observed under the deep respiratory regulation, whereas the reductions were much smaller in the absence of respiratory influence and under normal breathing. Specifically, MAP decreased by 2.20 mmHg without respiratory input, by 2.53 mmHg under normal respiration, and by 6.28 mmHg under deep respiration. The stepwise decline demonstrates that deep respiratory modulation enhances both the sensitivity and efficacy of PSNS regulation, thereby increasing the extent of BP recovery.

The observed improvement in regulatory performance may be associated with low-frequency ITP oscillations introduced by deep respiration. These oscillations can repeatedly shift the pressure signal into the high-slope region of the baroreflex sigmoid response function, particularly under conditions of elevated PSNS weighting. Increased gain within this region may amplify the PSNS contribution to the overall control signal σ, resulting in a greater reduction in MAP. While the simulations are consistent with this mechanism, the physiological processes underlying this effect require further empirical validation. Nonetheless, the model results suggest that following BP elevation induced by acute stress or psychiatric conditions, prolonged respiratory cycles or increased breathing depth may support ANS-mediated BP recovery.

## 6. Discussion

Although previous studies have investigated multisystem physiological interactions using experimental and computational approaches, autonomic regulatory mechanisms underlying interoceptive modulation—and their relationships with neurodevelopmental conditions—have not been explicitly quantified or systematically characterized. Within the proposed modeling framework, this study defined ANS functions as autonomic control modes and relative activity of each ANS branch. Differences of ANS regulation between TD and ASD individuals were also estimated via computational model integrating multiple physiological models.

### 6.1 Estimation of ANS functions in TD and ASD individuals

Compared with TD individuals, this study indicated the ASD group was predominantly distributed in regions characterized by concurrently high SNS and high PSNS relative activity across all three autonomic control modes. This pattern indicates that PSNS relative activity in the ASD group was not comparably reduced under external stress. Such a profile suggests blunted parasympathetic reactivity, in which PSNS relative activity cannot be flexibly adjusted in response to external demands, potentially constraining effective modulation of physiological signals and following behavioral regulation. Muscatello et al. [55] reported blunted PSNS responses in older children with ASD during peer interaction tasks, which were associated with altered stress responsivity and behavioral regulation. Similarly, Ward et al. [56] observed that higher PSNS relative activity was positively associated with ADHD symptom severity during short-term memory storage, suggesting that elevated or insufficiently modulated PSNS activity may contribute to reduced responsiveness to external cognitive or environmental demands. The present model may offer a computational account of these prior findings by characterizing autonomic regulation in terms of branch-specific relative activity, thereby providing an interpretable framework for understanding blunted parasympathetic reactivity in ASD.

In addition, the present findings indicate ASD individuals exhibit pronounced instability in autonomic regulation, as convergent SNS–PSNS weighting patterns render multiple autonomic control modes similarly capable of accounting for their ANS responses under identical external stimulation. ANS instability is associated with elevated interoceptive prediction errors transmitted to higher-order neural systems, as emphasized in theoretical accounts of interoceptive inference [23]. Such elevated prediction errors have been proposed to contribute to cognitive experiences marked by reduced confidence, diminished clarity, and a weakened sense of reality [24]. ANS instability reduces strategy specificity and limits the brain’s ability to correctly attribute bodily changes to internal or external causes [25]. This impaired attribution may further disrupt the integration of bodily signals with perceptual and contextual information, thereby destabilizing subjective experience, including emotional regulation and cognitive processes. Taken together, the observed autonomic instability, reduced interoceptive precision, elevated prediction error, and impaired attribution delineate a mechanistic sequence linking altered autonomic regulation to changes in cognitive experience. This physiological-to-cognitive framework provides a coherent interpretation of the empirical findings and supports the view that atypical autonomic dynamics may contribute to differences in perceptual and experiential processing in ASD.

We also systematically analyzed the HR trajectories as an example to discuss the determinants of variation directions and magnitudes of ANS-modulated interoceptive signals. Within the ANS regulatory framework, directions and magnitudes of interoceptive variation were determined by the sign and the absolute value of difference between baseline level and initial level. However, under specific combinations of SNS-PSNS relative activity, HR failed to exhibit a compensatory increase following the HUT stimulus and instead showed a further reduction, deviating from the typical physiological response observed in healthy individuals after postural transition. For example, the model predicted that ANS-modulated HR stabilized at a level lower than the regulatory target *μ* when both *α* and *β* were set to 1.0 [Fig 4]. It is worth noting that HR-suppressive responses emerging under high PSNS gain conditions within the model are not entirely absent in physiology, which delineate a potential pathological ANS regulatory phenotype. Multiple HUT studies in patients with vasovagal syncope have reported that, in a subset of individuals, HUT elicits abnormal cardioinhibitory response patterns characterized by abnormally enhanced PSNS activity and pronounced HR suppression [46]-[47]. In severe cases, this pathological phenotype can lead to marked bradycardia or transient cardiac asystole [48]. Accordingly, within a clinical context, HR decreases following HUT are generally interpreted as a manifestation of ANS dysregulation rather than normal compensatory regulation. The empirical HR distribution of the TD group on the simulated response surface [Fig 3] was concentrated in regions with high SNS and low PSNS weightings, where the model produced post-HUT HR elevations consistent with compensatory regulation. Although HR-suppressive patterns emerged in other SNS-PSNS relative activity regions, these did not overlap with the empirical HR of TD group. Overall, these findings demonstrate that the present model captures ANS regulatory patterns consistent with physiological expectations under healthy conditions while also explicitly identifying abnormal regulatory behaviors associated with ANS dysfunction within the broader parameter space. This capacity provides a structured computational framework for interpreting HR response mechanisms across distinct ANS regulatory states.

This study implemented unified baseline interoceptive signals as an ideal autonomic target to avoid masking abnormal initial SNS and PSNS states in the ASD group, thereby enabling fair comparisons of regulatory efficiency and deviations from the expected target between TD and ASD groups. Importantly, baseline interoceptive levels in the present framework represent ideal regulatory targets following stimulation, yet these targets may themselves be progressively elevated by sustained physiological or psychological stress. This interpretation aligns with allostatic load (AL) theory, which describes the cumulative physiological burden generated by repeated or chronic regulatory activation. Chronically elevated interoceptive states, such as persistently increased HR, may be incorporated into autonomic reference values, shifting the internal regulatory set point upward. Model simulations [Fig 5] illustrate that excessively elevated baselines restrict the effective regulatory range and yield blunted HR responses that lack typical compensatory dynamics. Although such set point shifts may support short-term stability, they reduce system sensitivity and limit effective autonomic compensation. Meanwhile, elevated allostatic load has been shown to be associated with reduced physical functioning, affective disturbances, and increased cognitive vulnerability [49],[50].

Accordingly, the unified baseline approach serves not only as a methodological requirement for TD–ASD comparisons, but also as a physiologically grounded framework for interpreting how sustained physiological burden may shape atypical ANS regulation and its cognitive implications.

### 6.2 Functional capabilities of the computational model

Previous ANS regulation models based on baroreceptor feedback designate BP as the primary regulatory target [42]-[44], while HR modulation is treated as a secondary mechanism to stabilize BP. Although HR components may be included structurally in such models, parameterization and optimization typically remain BP-centered, lacking a systematic framework in which HR is modeled as an independent output variable. This limitation reduces explanatory power when applied to physiological or psychological conditions characterized by asynchronous HR and BP responses, including disease states, autonomic dysfunction, or neurodivergent phenotypes. For example, Fukuda et al. [18] and Kollai et al. [19] reported that under severe hypoxia, both HR and BP increase to enhance oxygen exchange efficiency, whereas under mild hypoxia, BP increases while HR often remains stable, preserving hemodynamic balance. Differences in modulatory mechanisms between severe and mild hypoxia indicate that HR and BP are not always regulated synchronously and that the ANS may adopt differentiated control strategies depending on the physiological context. The present computational model enables independent modulation of HR and BP, which is therefore physiologically justified and a capability not supported by previous baroreflex-based models.

Importantly, the responses of the ANS branches (*n*_*s*_and *n*_*p*_) and their relative activity weights (*α* and *β*) play distinct yet complementary roles. The response levels of the SNS and PSNS represent the immediate activation of the ANS in response to incoming interoceptive signals (current HR and BP levels), whereas the relative activity parameters *α* and *β* scale the peripheral effectiveness of SNS and PSNS outputs on target organs, including the heart and blood vessels. ANS function is typically inferred from aggregate cardiovascular outputs in experimental or clinical settings, whereas internal regulatory components cannot be independently isolated or systematically manipulated. By contrast, the present model allows controlled variation of parameters associated with distinct regulatory levels, enabling separate examination of neural activation patterns and their effectiveness in cardiovascular regulation. This structured decomposition therefore provides a mechanistic framework for exploring how combinations of autonomic responsiveness and modulation efficacy contribute to interoceptive dynamics, thereby facilitating more detailed comparisons of autonomic regulation between TD and ASD individuals than is possible using empirical measurements alone.

In addition to simulating HR and BP elevations in response to external stimulation, the proposed model can also predict recovery trajectories following stimulus cessation. As a practical application, a respiratory module was introduced to examine how distinct breathing patterns modulate BP recovery. This module simulates normal and deep breathing by adjusting the frequency and amplitude of ITP fluctuations, manifesting in the BP waveform as lower-frequency, longer-period components relative to the cardiac cycle. Model simulations indicated that, in the absence of breathing and normal breathing, elevated PSNS activity alone was insufficient to induce substantial BP reduction when SNS activity remained high. In contrast, when deep breathing at a rate of 9 bpm was introduced, PSNS contributions to BP recovery were markedly enhanced even under elevated SNS tone, suggesting that deep breathing facilitates PSNS-driven modulation. Consistent with these findings, prior studies have shown that resonance-frequency breathing reduces blood pressure, enhances heart rate variability, and shifts autonomic balance toward increased PSNS and reduced SNS activity, alongside improvements in stress, anxiety, and cognitive function [51]-[52].

### 6.3 Limitations and future work

The computational model explicitly distinguishes multiple regulatory parameters across different functional levels, only a restricted subset, however, of the parameter space was systematically explored. In particular, the present analysis intentionally adopted a unified baseline (*μ*) to ensure comparability of autonomic regulatory performance between TD and ASD individuals, but did not examine how group-specific baseline settings or variations in autonomic response sensitivity interact with afferent activation and control modes. Moreover, the response slopes of the sympathetic and parasympathetic branches (*u*_*s*_ and *u*_*p*_), which determine the sensitivity of autonomic responses to deviations from baseline, were not independently manipulated. Future studies must extend the current framework by systematically varying baseline values and response slopes to examine how baseline shifts, response sensitivity, and relative activity weighting jointly shape interoceptive regulation. These extensions will enable a more comprehensive characterization of adaptive versus maladaptive autonomic strategies and clarify conditions under which elevated baselines reflect allostatic accommodation rather than regulatory efficiency and enhance the model’s ability to capture individual and group-level heterogeneity in autonomic control.

Although the proposed model explicitly distinguishes multiple regulatory parameters across functional levels, its validation was primarily based on a single empirical dataset from a prior study. This reliance on one dataset limits the generalizability of the findings to some extent and constrains the extent to which inter-individual and group-level heterogeneity in autonomic regulation can be fully characterized. Future work should therefore evaluate the model across independent datasets and experimental paradigms to assess its robustness and to determine whether the observed regulatory patterns generalize across populations, task contexts, and physiological conditions. The present model currently focuses on cardiovascular interoceptive signals, which limits its scope to a single physiological domain. Future work should extend this framework by integrating respiratory, electrodermal, and gastrointestinal systems to investigate multimodal ANS regulation and to assess whether similar nonlinear regulatory principles generalize across interoceptive domains, particularly in neurodivergent populations.

Building on the present modeling framework, several extensions will be important for further advancing the quantitative characterization of autonomic regulation. Although respiratory-related PSNS gains contributing to BP restoration could be observed by incorporating deep respiration, the underlying mechanisms remain to be further elucidated. In addition, there are no direct interaction mechanisms between the ANS and respiratory systems implemented in current framework, such that the gradual decline in the overall blood pressure level over a long period of time cannot be effectively replicated or explained by adjusting the function of the ANS or the breathing pattern. These limitations motivate future development of a more comprehensive modeling framework to elucidate respiratory–ANS interactions and to support the design of physiologically grounded, non-pharmacological self-regulation strategies, particularly for neurodivergent individuals with ANS dysfunction. More broadly, incorporating additional new interaction mechanisms would enable the integrated analysis and comparison of multiple interoceptive signals—such as cardiovascular, respiratory, electrodermal, and gastrointestinal dynamics—within a unified ANS regulation framework, facilitating the development of a multimodal simulation platform with stronger system-coupling properties and supporting ANS function assessment, interoceptive regulatory mechanism analysis, and intervention and prediction across a wider range of neurodivergent populations.

## 7. Conclusions

This study developed a closed-loop computational framework integrating the CVS, RS, and ANS to systematically estimate autonomic regulatory functions underlying interoceptive modulation in TD and ASD individuals. By explicitly parameterizing autonomic control modes and branch-specific relative activities, the framework enables quantitative comparison of autonomic regulatory strategies across neurodevelopmental conditions, extending beyond descriptive characterization of autonomic dysfunction. HR and BP responses to the head-up tilt (HUT) test were simulated, and regulatory surfaces were compared with experimental HR and BP data from TD and ASD groups. TD individuals exhibited differentiated SNS– PSNS coordination patterns across control modes, whereas ASD individuals demonstrated a greater convergence of relative SNS–PSNS activity, suggesting reduced flexibility and potential instability in autonomic control strategies under the same stimulus. Analysis of empirical HR and BP distributions on simulated responses surfaces following the HUT test indicates that the coupled reciprocal control mode demonstrates greater capacity to capture intrinsic autonomic regulatory processes in TD individuals. Under this mode, simulated SNS–PSNS relative activity patterns aligned most closely with expected physiological responses to postural challenge. In contrast, although the majority of ASD observations were likewise located within regions of elevated SNS activity, empirical data extended toward areas associated with relatively higher PSNS weighting on the simulated regulatory surfaces. This distributional shift suggests that increased parasympathetic involvement may limit effective ANS-mediated interoceptive adjustment in ASD, thereby reducing the adaptability of cardiovascular regulation under external stimulation. Simulations under the coupled reciprocal mode were performed under normal respiration, deep respiration, and absence of respiration to evaluate their effects on mean arterial pressure (MAP) across varying SNS–PSNS activity combinations. Incorporation of deep respiration enhanced PSNS effectiveness in lowering MAP during BP recovery, particularly under conditions of over-activity of SNS. Collectively, these findings establish a mechanistic, state-space framework for characterizing autonomic coordination by linking measurable interoceptive signals to their underlying control architectures. By enabling quantitative inference of regulatory stability in TD and ASD populations and identifying respiration as a model-based modulatory factor in cardiovascular recovery, the proposed approach provides a generalizable computational strategy for probing autonomic regulation across neurodevelopmental conditions.

